# Molecular Basis of High-light Adaptation in Cyanobacteria and Cyanophages through the D1/D2 subunits of Photosystem II

**DOI:** 10.64898/2026.07.15.738324

**Authors:** Rulong Ma, Ruonan Wu, Greg Morrison

**Affiliations:** Department of Physics, University of Houston, Houston, TX 77204, USA; Center for Theoretical Biological Physics, Rice University, Houston, TX 77005, USA; Integrated Discovery Sciences Directorate, Pacific Northwest National Laboratory, Richland, WA, USA

**Keywords:** Photosystem II, High-light adaptation, *Prochlorococcus*, Cyanophages, ROS, Protein evolution

## Abstract

High-light (HL) stress imposes substantial pressure on oxygenic photosynthesis by promoting the formation of reactive oxygen species (ROS) and photodamage in Photosystem II (PSII), particularly within its reaction-center core subunits D1 and D2. *Prochlorococcus* and *Synechococcus,* along with their cyanophages, dominate oligotrophic oceans and experience persistent HL exposure, yet the molecular basis of PSII adaptation in these systems remains poorly understood. In this manuscript, we integrate large-scale phylogenetic analysis with residue-level sequence comparison and AlphaFold3-based structural prediction to investigate D1/D2 across *Prochlorococcus* ecotypes, *Synechococcus*, and their associated cyanophages. *Prochlorococcus* species are known to form ecotypes based on light conditions, and the D1 and D2 phylogenies showed consistent organization across our phylogenetic trees. *Synechococcus* does not fall into the same light-based ecotypes, but we found that D1 and D2 isoforms still form meaningful clusters for these species. Cyanophage-encoded D1 and D2 sequences did not form separate viral clades, but instead were embedded within host-associated diversity and showed closer affinity to HL-adapted *Prochlorococcus* than to low-light-adapted (LL) lineages. Mapping ecotype- and phage-associated substitutions onto predicted D1/D2 structures, with reference to experimental PSII structures, suggested that some LL-to-HL variants in *Prochlorococcus* may enhance resistance to ROS-associated damage. Some LL-to-HL substitutions increase methionine and cysteine enrichment near redox-active cofactors, potentially improving redox buffering and ROS scavenging. In contrast, others may favor more efficient electron transfer, reduce charge recombination, and thereby limit ROS production. These results support a model in which PSII adaptation in cyanobacteria is shaped by fine-scale protein-level tuning linked to ecological diversification and cyanophage-host coevolution, with viral-host interactions contributing to the evolutionary optimization of oxygenic photosynthesis under HL stress.

## 1. Introduction

Marine picocyanobacteria of the genera *Prochlorococcus* and *Synechococcus* dominate oligotrophic oceans and together account for approximately one quarter of global marine primary production, forming the energetic foundation of pelagic food webs and exerting a major influence on the global carbon cycle (*1*). Among these taxa, *Prochlorococcus* exhibits extraordinary ecological and evolutionary diversification, having radiated into multiple genetically and physiologically distinct ecotypes that occupy sharply defined niches along vertical gradients of light, temperature, and nutrient availability (*2*, *3*).

*Prochlorococcus* ecotypes are broadly divided into low-light (LL) and HL (HL)-adapted lineages. LL-adapted clades are distributed progressively deeper in the water column, with LLI occurring near the nutricline, LLII/III occupying slightly deeper strata, and LLIV residing near the lower boundary of the euphotic zone (*3*). Recently described LLVII and LLVIII lineages further expand this diversity and are adapted to extremely low irradiance, exhibiting distinct pigmentation patterns and light-harvesting gene repertoires that distinguish them from classical LL ecotypes (*4*). In contrast, HL-adapted *Prochlorococcus* dominate surface waters and are subdivided according to iron availability and temperature, including low-iron–adapted clades (HLLIII/IV) and high-iron–adapted clades that further separate into low-temperature (HLI) and high-temperature (HLII) groups (*5–9*). This fine-scale correspondence between phylogeny and environment suggests that light intensity is a major ecological factor associated with *Prochlorococcus* diversification. *Synechococcus* does not fall into similar HL and LL ecotypes (as discussed below).

In cyanobacteria, light energy is captured and converted into chemical energy by Photosystem II (PSII), a large multisubunit pigment–protein complex embedded in the thylakoid membrane. PSII catalyzes the light-driven oxidation of water at the oxygen-evolving complex (OEC), generating molecular oxygen, protons, and electrons that enter the photosynthetic electron transport chain (*10*). The reaction center of PSII is formed by the D1 (encoded by *psbA*) and D2 (encoded by *psbD*) proteins, which coordinate chlorophyll *a* molecules (P_D1_/P_D2_ and Chl_D1_/Chl_D2_), pheophytin (Pheo), plastoquinones (Q_A_ and Q_B_), and redox-active amino acid residues that collectively enable directional charge separation and electron transfer.

Despite its central role in oxygenic photosynthesis, PSII is vulnerable to photodamage under HL conditions, when the absorbed excitation energy may exceed the capacity for photochemistry or regulated dissipation (*11*). Excess excitation promotes the formation of reactive oxygen species (ROS) through multiple pathways, including charge recombination that generates triplet chlorophyll, which subsequently produces singlet oxygen; direct reactions of excited pheophytin with molecular oxygen; and electron leakage from reduced plastoquinone intermediates to oxygen, producing superoxide radicals (*11–15*). These ROS are more likely to damage the PSII reaction center, with the D1 subunit being the primary target of oxidative injury, necessitating continual repair or replacement to sustain photosynthetic function.

In plants and algae, photodamage is mitigated by multilayered defense strategies that include chloroplast avoidance movements, accumulation of photoprotective pigments, non-photochemical quenching, cyclic electron flow, photorespiration, antioxidant networks, and highly regulated PSII repair cycles (*16–18*). In contrast, marine picocyanobacteria are free-floating unicellular organisms with limited capacity for spatial or behavioral light avoidance. As a result, HL imposes a persistent selective pressure on its photosynthetic machinery. In *Prochlorococcus*, this pressure is further intensified by extreme genome streamlining. Each strain encodes only a single copy of *psbA* and *psbD*, eliminating paralog redundancy observed in other genera and making protein-level adaptations within D1 and D2 especially critical for PSII robustness under HL stress. By comparison, many *Synechococcus* strains encode multiple D1 isoforms, most notably D1.1 and D1.2, which are differentially expressed under normal and HL conditions, respectively, thereby providing a transcriptionally regulated mechanism for mitigating photodamage (*19*). *Prochlorococcus*, lacking this flexibility, may rely more on constitutive sequence features of its PSII core proteins to withstand HL stress, making it a good model for identifying protein-level mechanisms of PSII stress tolerance.

Adding further complexity, numerous cyanophages infecting *Prochlorococcus* and *Synechococcus* encode auxiliary metabolic genes (AMGs) involved in photosynthesis, including *psbA* (encoding the D1 subunit) and, in a substantial subset, *psbD* (encoding the D2 subunit) (*20–23*). Phage-encoded D1 and D2 proteins are expressed during infection. They can sustain or even enhance host photochemical activity under HL conditions, suggesting that viral PSII components may experience selective pressure for photodamage resistance (*12*, *20*, *24*). However, the molecular basis by which phage-encoded reaction-center proteins contribute to HL tolerance and how these adaptations relate to host ecotype-specific structural features remain poorly understood.

Previous evolutionary studies have classified cyanobacterial D1 proteins into multiple functional types (G0–G4, D1^INT^, and D1^FR^), reflecting differences in oxygen-evolving capacity, redox tuning, and spectral adaptation (*16*, *17*). The G0-type D1 from *Gloeobacter kilaueensis* JS1, which lacks key ligands for the oxygen-evolving complex (OEC), is considered the most ancestral form and has been widely used as an evolutionary outgroup. While this framework has yielded important insights into early PSII evolution, most classifications have relied on gene-level phylogenies and have not explicitly addressed how protein-level sequence variation maps onto PSII structure to confer HL adaptation within ecologically structured marine cyanobacteria and their phages.

Although large-scale surveys have established that cyanophage and host PSII genes are evolutionarily intertwined through horizontal transfer, recombination, and lineage sorting, these analyses have remained largely sequence-centric, with limited structural interpretation-particularly for D2, which is less frequently encoded by phages and consequently less studied than D1 (*20–23*). Recent advances in protein structure prediction (e.g., AlphaFold) now enable structure-aware evolutionary analysis of PSII core subunits beyond the limited set of experimentally resolved complexes, opening new opportunities to test mechanistic hypotheses about photodamage resistance in an explicitly three-dimensional framework (*25*).

Here, we integrate ecotype-resolved phylogenetics, residue-level sequence comparison, and AlphaFold3-based structural prediction to investigate how the D1 and D2 proteins of *Prochlorococcus*, *Synechococcus*, and their cyanophages differ between HL and LL adaptations. By mapping ecotype- and phage-specific substitutions onto PSII structural contexts, we identify multiple protein-level strategies-including oxidation-resistant substitutions, ROS-suppressing redox modulation, and redox-buffering residue distributions-that may contribute to PSII robustness under HL stress. Our results indicate that D1/D2 sequence variation in marine picocyanobacteria is ecotype-structured and closely associated with the genetic pool of cyanophages, highlighting a potential viral contribution to the molecular diversification of oxygenic photosynthesis.

The paper is organized as follows: In Sec. 2, we describe the dataset of *Prochlorococcus, Synechococcus,* phage, and other sequences of D1 and D2 PSII subunits used in this paper. We describe the phylogenetic analysis of the D1 and D2 subunits and the implications of each in Sec. 3.1 and 3.2, respectively. A comparison of these trees, including potential evolutionary implications, is found in Sec. 3.3. In Sec 3.4, specific substitutions that distinguish the clades corresponding to the HL and LL ecotypes in *Prochlorococcus* are identified. These substitutions are classified based on structural modeling into Oxidation-Resistant Substitutions (ORSs), ROS-Suppressing Substitutions (RSSs), or Redox Buffering Substitutions (OBSs) in Sec 3.5, 3.6, and 3.7, respectively. Finally, in Sec. 4 we discuss the overall implications of the combined phylogenetic and structural analysis presented here.

## 2. Methods

### 2.1. Dataset Construction for D1 and D2 Phylogenetic Analyses

To investigate the evolution of the PSII D1 and D2 proteins, we collected their sequences and constructed datasets for phylogenetic analyses. Because this study focuses on *Synechococcus*, *Prochlorococcus*, and their associated cyanophages, we curated an extensive collection of D1/D2 sequences from these groups. To place these lineages in a broader evolutionary context, representative taxa from other cyanobacteria, algae, and higher plants that had previously been analyzed in comparative PSII studies were also included (*26*, *27*). All protein sequences were retrieved from GenBank (ncbi.nlm.nih.gov) or UniProt (uniprot.org).

For *Synechococcus*, we collected D1 and D2 protein sequences from 242 *Synechococcus* strains (Table 1). Among these, we identified 194 distinct D1 homologs derived from 166 strains, reflecting the widespread presence of multiple *psbA* paralogs within many *Synechococcus* genomes and the resulting production of distinct D1 protein types, most notably D1.1 and D1.2. We also recovered 102 D2 homologs from 98 *Synechococcus* strains. To ensure phylogenetic robustness, only full-length D1/D2 sequences were retained for D1/D2 tree reconstruction.

**Table 1.** Summary of organisms and corresponding numbers of D1 and D2 protein sequences analyzed.

| **Organisms** | **Numbers** | **D1 protein** | **D2 protein** |
| --- | --- | --- | --- |
| *Prochlorococcus* HL strains | 23 | 23 | 23 |
| *Prochlorococcus* LL strains | 38 | 38 | 38 |
| *Prochlorococcus* phages | 23 | 23 | 8 |
| *Synechococcus* strains | 242 | 194 (166) * | 102 (98) * |
| *Synechococcus* phages | 104 | 104 | 72 |
| Representative Cyanobacteria | 12 | 32 | 14 |
| Algae | 5 | 5 | 5 |
| Plants | 7 | 7 | 7 |
| Total | 454 | 426 | 269 |

**Note\*:** For entries formatted as $X (Y)$ , $X$ denotes the number of protein sequences, whereas $Y$ denotes the number of cyanobacterial strains in which those sequences are present.

Because this study focuses on light-adapted *Prochlorococcus* ecotypes, we selected 23 high-light (HL)-adapted and 38 low-light (LL)-adapted strains with well-defined ecotype classifications (Table S1). Strain information was obtained from the Chisholm Lab *Prochlorococcus* stock culture resource (Table S1). These strains represent the major Prochlorococcus sub-ecotypes HLI, HLII, LLI, LLII/III, LLIV, LLVII, and LLVIII. They were collected from diverse marine environments, including the Mediterranean Sea, Arabian Sea, and the North and South Pacific and Atlantic Oceans, across depths ranging from the surface to 200 m. Each *Prochlorococcus* genome encodes a single homolog of both the D1 and D2 proteins, simplifying evolutionary interpretation by avoiding paralogous ambiguity. Protein sequences were retrieved from GenBank or UniProt.

Cyanophage sequences were also extensively sampled. We identified 23 *Prochlorococcus* cyanophages that encode the *psbA* gene, of which 8 also encode *psbD,* as reported in a previous paper (*23*) or in GenBank (Table 1, S2). In addition, 104 *Synechococcus* cyanophages encoding *psbA* were included, 72 of which also carry *psbD*. In total, 127 cyanophages were analyzed in this study, substantially expanding upon previous phylogenetic analyses that examined fewer than 70 cyanophages carrying viral photosynthesis genes (*23*). Further details are provided in the Supplementary Materials, including a list of cyanobacteria–cyanophage host–virus interactions reported in the literature (Table S2).

Five algal species with experimentally resolved PSII structures were included to anchor sequence-based analyses to structural information (Table 1). These taxa include *Cyanidium caldarium* (red alga; PDB: 4YUU), *Chlorella ohadii* (green alga; PDB: 8BD3), *Chlamydomonas reinhardtii* (green alga; PDB: 8KDE), *Chaetoceros neogracilis* (diatom; PDB: 7VD5), and *Rhodomonas salina* (cryptophyte; PDB: 8XLP). Corresponding D1 and D2 protein sequences were extracted directly from their experimentally determined PSII structures. In addition, seven plant species analyzed in previous studies were included to represent higher-plant PSII evolution (*27*).

Finally, 12 representative cyanobacterial species were selected to comprehensively capture the known diversity of cyanobacterial D1 proteins (Table 1). Previous studies have classified cyanobacterial D1 proteins into several distinct functional groups, including G0–G4, D1^INT^ (intermediate), and D1^FR^ (far-red–adapted). The G0-type D1 from *Gloeobacter kilaueensis* JS1, which lacks a clearly defined photosynthetic function, is thought to represent the most ancestral D1 form and has been used as an outgroup in prior evolutionary analyses (*16*, *26*).

Several cyanobacteria—*Chlorogloeopsis fritschii* PCC 6912, *Fischerella* sp. JSC-11, and *Fischerella* sp. PCC 9605—were included because each genome encodes multiple *psbA* paralogs, which collectively span the G1, G2, G4, D1^INT^, and D1^FR^ D1 functional groups; likewise, *Cyanothece* sp. PCC 7425, *Planktothrix serta* PCC 8927, and *Thermosynechococcus elongatus* BP-1 were selected because each genome harbors multiple *psbA* paralogs covering the G3 and G4 groups.

In addition, D1 and D2 sequences from cyanobacterial PSII complexes with experimentally determined structures were incorporated, specifically from *Synechocystis* sp. PCC 6803 (PDB: 7N8O), *Thermosynechococcus vestitus* BP-1 (PDB: 9EVX), *Thermostichus vulcanus* (PDB: 8IR5), *Synechococcus* sp. PCC 7335 (PDB: 8EQM), and *Acaryochloris marina* MBIC11017 (PDB: 7YMI). In summary, the D1 protein sequences from these 12 representative cyanobacteria collectively encompass all currently recognized D1 functional types, providing a robust evolutionary framework for comparative analysis of PSII reaction-center proteins.

### 2.2. Phylogenetic analysis of D1/D2 proteins

Separate phylogenetic trees were constructed for the D1 and D2 protein datasets, which contained 426 and 269 sequences, respectively, as described above. Duplicate protein sequences from the same organisms were removed from both datasets. Because some phage-encoded D1 and D2 proteins contained extended N-terminal regions, these extensions were trimmed before multiple sequence alignment. Since the N-terminal region varies among organisms, most-but not all-start-codon methionines were aligned across protein sequences. Multiple sequence alignments were performed independently for each dataset using MUSCLE with default parameters (gap opening penalty = -2.9, hydrophobicity multiplier = 1.2, clustering method = UPGMA, minimum diagonal length = 24) in MEGA v12.1 (*28*).

The most appropriate amino acid substitution model for each dataset was identified using the Models module in MEGA v12.1. For both D1 and D2, the best-fitting model was the Le and Gascuel (LG) model with gamma-distributed rate heterogeneity and a proportion of invariant sites, using five discrete categories (LG+G+I). Maximum-likelihood phylogenetic trees were reconstructed in MEGA v12.1. Statistical support for the inferred topologies was evaluated using 1,000 bootstrap replicates, and bootstrap values were assigned to the consensus trees. Gaps and missing data were treated using the Partial Deletion method with a 95% site-coverage cutoff. Heuristic ML searches were performed using the Subtree-Pruning-Regrafting (SPR, level 3) algorithm, with starting trees generated automatically by the Neighbor-Joining (NJ) or BioNJ methods based on pairwise distance matrices. Final trees were visualized and annotated in iTOL, and the resulting alignments were examined in Jalview v2.11.5.0 (*29*).

### 2.3. Protein structure prediction, cofactor mapping, and structural analysis

Because no experimentally resolved PSII complex structure is currently available for *Prochlorococcus* strains, the structural complex of the D1, D2, and F subunits of *Prochlorococcus* strains MED4 and MIT9313 was predicted using AlphaFold3 without explicitly including cofactors (*30*). The input protein sequences were obtained from GenBank or UniProt. The model with the highest confidence score was used for the subsequent analyses. To approximate the positions of PSII cofactors in the predicted *Prochlorococcus* complex, the predicted D1/D2/F structures were structurally aligned to the experimentally resolved PSII complex from *Synechocystis* sp. PCC 6803 (PDB ID: 7N8O). Cofactor coordinates from 7N8O were then used as structural references to map the corresponding cofactor-binding regions in the predicted *Prochlorococcus* D1/D2/F complex. Structural alignments and visualization of these subunits were performed using VMD (version 1.9.4) (*31*).

## 3. Results

### 3.1. Phylogenetic analysis of PSII D1 (*psbA*) proteins

Using 426 D1 protein sequences sampled from 454 organisms or cyanophages (Table 1), we reconstructed a maximum-likelihood (ML) phylogeny of the PSII reaction-center D1 subunit. The tree was rooted with the G0-type D1 from *Gloeobacter kilaueensis* JS1 as an outgroup, consistent with its established position as one of the most ancestral D1 forms. The tree is shown in Fig 1, with colors indicating the specific groups of organisms or viruses considered.

**Figure 1.**
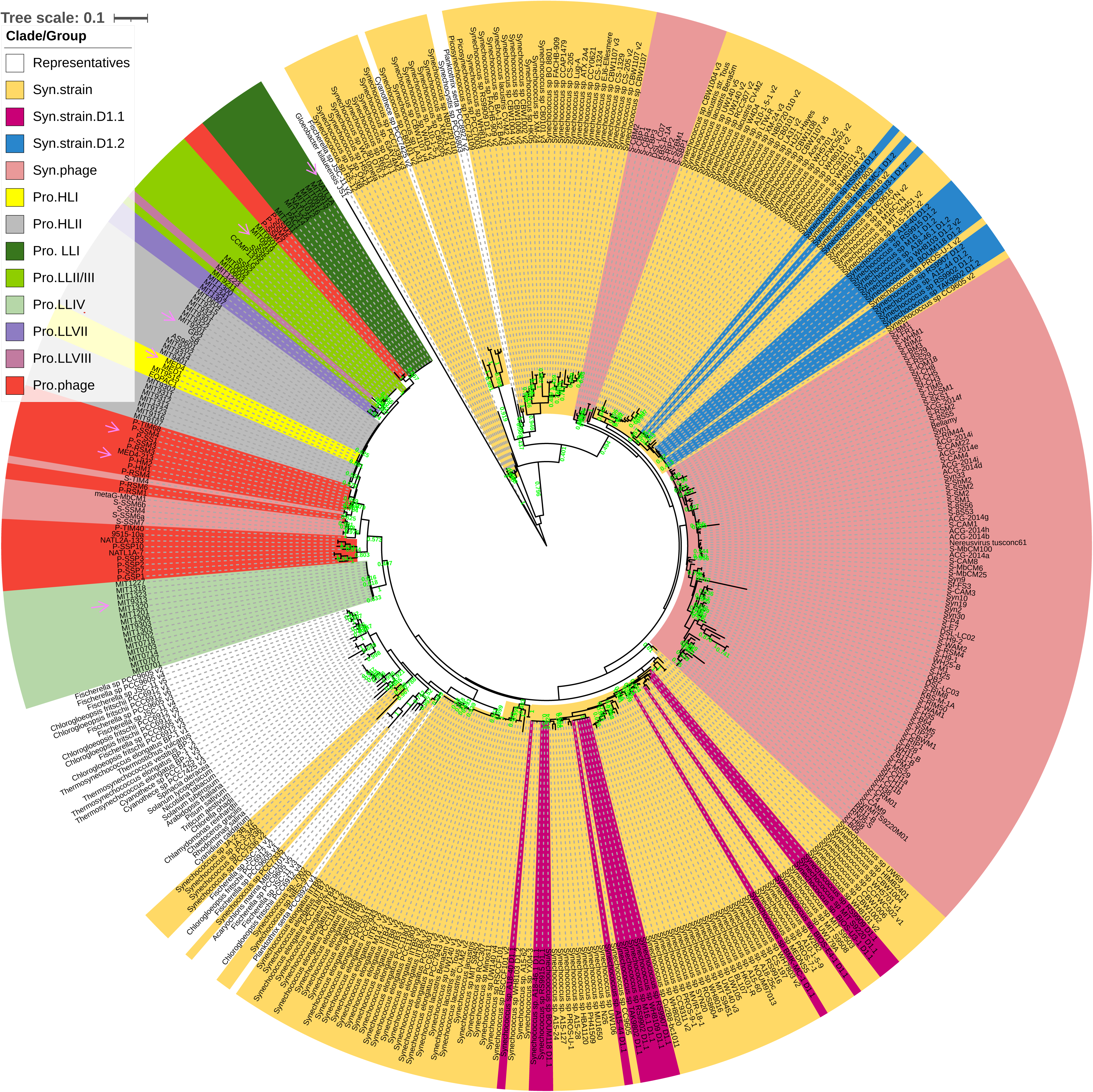
Rooted maximum-likelihood phylogeny of D1 proteins, with the atypical G0-type D1 from *Gloeobacter kilaueensis* JS1 used as the outgroup. In the tree, *Prochlorococcus* sequences are denoted as “**Pro**,” and *Synechococcus* sequences are denoted as “**Syn**.” Distinct clades corresponding to *Prochlorococcus*, *Synechococcus*, and their associated cyanophages are highlighted in different colors. The G0 sequence from *Gloeobacter kilaueensis* JS1, together with representative **G1, G2, G3, and G4 D1 proteins** from cyanobacteria, algae, and plants, is shown in white. Bootstrap support values ≥0.4 are shown at internal nodes. Branch lengths are proportional to the inferred number of amino acid substitutions per site; the scale bar represents 0.1 substitutions per site. Pink arrows indicate representative strains and phage sequences analyzed below in the phylogenetic tree.

#### 3.1.1. Global D1 backbone separates host lineages from viral radiations

Across the whole phylogeny, D1 sequences segregate into a coherent host-derived backbone that includes (i) representative cyanobacteria spanning recognized cyanobacterial D1 functional diversity and (ii) eukaryotic algal/plant D1 proteins, which together provide a broad evolutionary context for the marine picocyanobacteria. In the circular tree representation, these reference lineages occupy a distinct “representatives” sector that anchors the interpretation of the densely sampled marine clades.

Superimposed on this host backbone, the dataset reveals extensive viral expansion: D1 sequences encoded by cyanophages form large, coherent radiations that are visually and topologically separated into *Synechococcus* phage and *Prochlorococcus* phage partitions. This organization is consistent with long-term acquisition and diversification of viral *psbA* genes within host-associated gene pools, while still permitting recurrent exchange among related marine phototroph communities.

#### 3.1.2. *Prochlorococcus* D1 clades follow ecotype structure

Because each *Prochlorococcus* genome encodes a single D1 (*psbA*) homolog, the *Prochlorococcus* portion of the D1 phylogeny is free of paralog ambiguity and partitions the tree into known ecotypes. In our dataset, D1 sequences from 23 HL and 38 LL-adapted *Prochlorococcus* strains span all major ecotype clades: HLI, HLII, LLI, LLII/III, LLIV, LLVII, and LLVIII (Table S1). These ecotypes form distinct and recognizable partitions in the D1 phylogeny (colored wedges in Fig. 1), indicating that D1 diversification in *Prochlorococcus* is likely shaped by long-term lineage splitting associated with light-adapted niche specialization.

The *Prochlorococcus*-centered region of the D1 phylogeny includes both host D1 sequences and *Prochlorococcus*-infecting cyanophage-encoded D1 sequences. Although the phage sequences form a tight subcluster, they fall within or adjacent to the broader Prochlorococcus-associated clades without forming a deeply separated viral lineage elsewhere in the tree (Fig. 1). We will refer to this as the *Prochlorococcus* clade in this paper. The smallest clade containing all *Prochlorococcus* D1 sequences comprises 61 host-derived D1 proteins and 29 cyanophage D1 proteins (total n = 90). The basal node defining this *Prochlorococcus* clade is supported by moderate bootstrap support (0.407), consistent with numerous short internal branches and/or recombination among closely related host and viral D1 lineages. Notably, cyanophage D1 sequences are embedded within host-associated diversity, rather than forming a distinct viral outgroup. This grouping is consistent with a model in which viral *psbA* evolution is tightly coupled to host D1 backgrounds, potentially through recurrent gene acquisition followed by sequence homogenization. However, this phylogenetic analysis cannot distinguish coevolution from recent host acquisition, recombination, or shared ancestry. More in-depth recombination analysis is needed to test the inference conclusively, but is beyond the scope of this work.

Several *Synechococcus* cyanophages, including S-TIM4, S-SSM6b, S-SSM4, S-SSM6a, and S-SSM7, also cluster within the *Prochlorococcus*-centered D1 clade. This placement is consistent with phage D1 sharing recent ancestry or genetic exchange with *Prochlorococcus*-like hosts. This may be potentially explained by horizontal acquisition of *psbA* across the host-associated gene pool. However, recombination analysis and host-range data would be required to determine the direction and mechanism of gene transfer. In addition, the metagenome-derived cyanophage metaG-MbCM1, for which no host has been experimentally identified, also falls within the *Prochlorococcus*-centered clade, suggesting *Prochlorococcus* as a potential host lineage.

Within the Prochlorococcus D1 clade, P-SSM2, P-SSM5, and P-SSM7 form a distinct cyanophage subclade positioned near the LLI ecotype. Although the larger phage and LLI assemblage has low bootstrap support (0.354), this placement is consistent with available host-range data: P-SSM2, P-SSM5, and P-SSM7 infect NATL1A and NATL2A (Table S2), both LLI Prochlorococcus strains. Thus, while the fine-scale phylogenetic placement should be treated cautiously, the combined tree topology and host-range data support an LLI-associated host background for these phage D1 genes.

#### 3.1.3. *Synechococcus* D1 phylogeny highlights extensive *psbA* paralogy and isoform diversification

In contrast to *Prochlorococcus*, the *Synechococcus* component of the D1 dataset exhibits strong paralog-driven expansion. Across 242 *Synechococcus* strains (Table 1), we identified 194 D1 homologs from 166 strains, reflecting the prevalence of multiple *psbA* paralogs per genome and the production of distinct D1 protein types, most prominently D1.1 and D1.2. This isoform structure is explicitly annotated in the tree as separate partitions for *Syn*. strain D1.1 and *Syn*. strain D1.2, embedded within the broader *Synechococcus* strain radiation. The topology indicates that D1.1 and D1.2 do not simply form a single clean “two-clade split” across all *Synechococcus*. Instead, isoform-bearing lineages recur throughout the broader diversity of *Synechococcus*.

#### 3.1.4. Cyanophage D1 sequences form major host-associated pools

Cyanophages comprise a substantial fraction of our D1 dataset: 23 *Prochlorococcus-infecting* phages and 104 *Synechococcus-infecting* phages encode *psbA* (Table 1). In the D1 tree, these sequences occupy large, visually distinct sectors labeled Pro. phage and Syn. phage, respectively. The clear partitioning of phage D1 sequences by host association supports the existence of host-structured viral *psbA* reservoirs, consistent with frequent phage–host gene exchange and subsequent diversification within ecological networks dominated by *Prochlorococcus* and *Synechococcus*. Importantly, the large size of the phage clade relative to the host reference groups may imply viral D1 evolution is not simply a passive reflection of host phylogeny. Instead, it may suggest recurrent acquisition of psbA, rapid sequence diversification, and recombination may together generate the observed phage partitioning.

Overall, the D1 phylogeny demonstrates three prominent patterns: (i) a deep host-derived backbone rooted by ancestral *Gloeobacter kilaueensis* G0-type D1, (ii) ecotype-structured diversification in single-copy *Prochlorococcus* D1, and (iii) extensive *psbA* paralogy and isoform diversification in *Synechococcus* coupled to large, host-associated cyanophage D1 radiations. Together, these results suggest a framework for subsequent comparative analyses of D1/D2 diversification, viral photosynthesis gene repertoires, and light-adaptation trajectories across marine picocyanobacteria and their phages.

### 3.2. Phylogenetic analysis of PSII D2 (*psbD*) proteins

We next reconstructed the phylogeny of the PSII reaction-center D2 (*psbD*) subunit using an unrooted maximum-likelihood (ML) tree (Fig. 2). The D2 dataset comprises 269 sequences, including marine picocyanobacteria (*Prochlorococcus* and *Synechococcus*), cyanophages encoding *psbD*, and representative cyanobacteria/algae/plants used to anchor deep PSII evolution. The resulting unrooted topology is shown with branch lengths scaled to substitutions per site (tree scale = 0.05) and annotated by major biological partitions.

**Figure 2.**
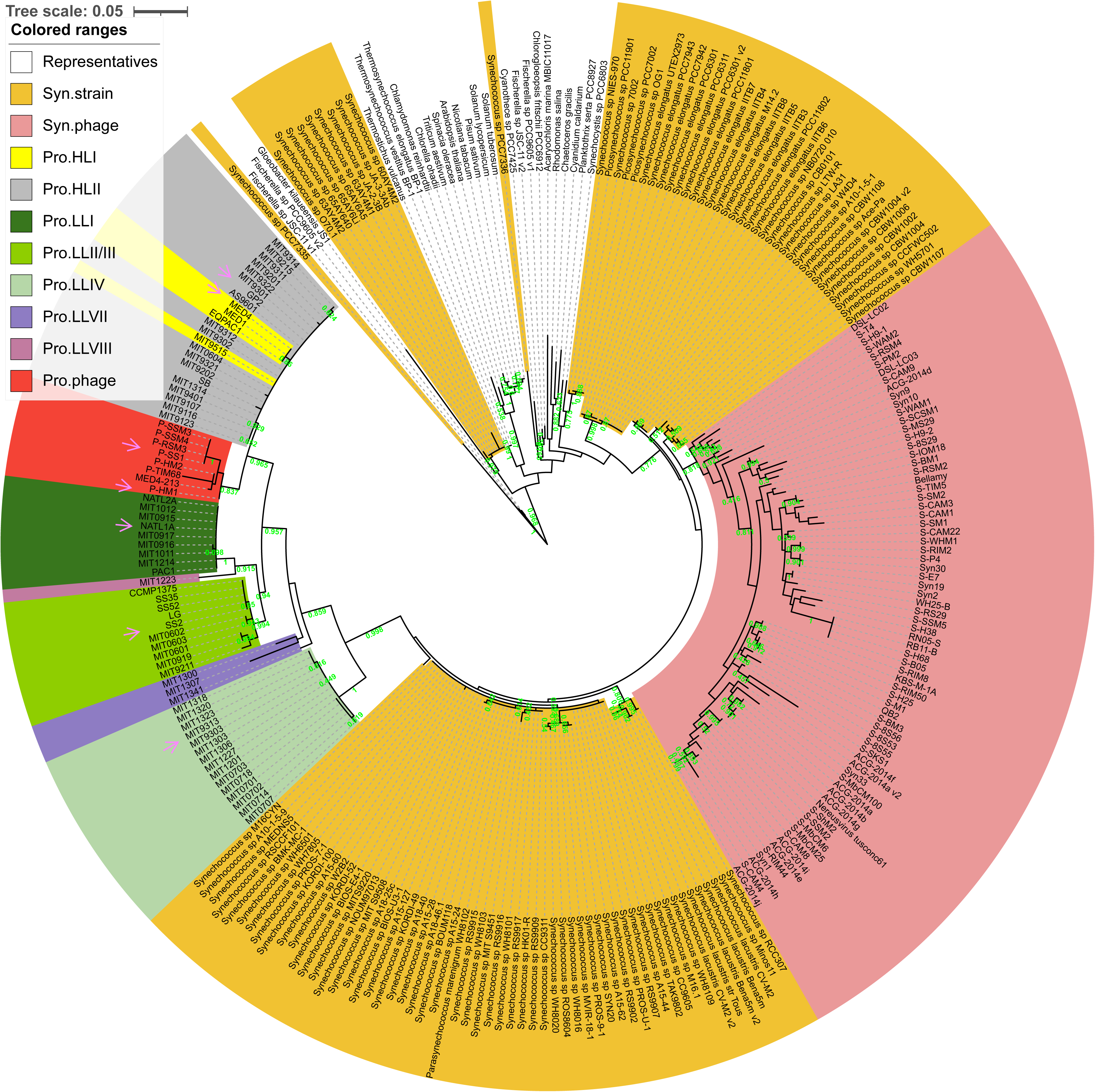
Unrooted maximum-likelihood phylogeny of D2 proteins. In the tree, *Prochlorococcus* sequences are denoted as “**Pro**,” and *Synechococcus* sequences are denoted as “**Syn**.” Distinct clades corresponding to *Prochlorococcus*, *Synechococcus*, and their associated cyanophages are highlighted in different colors. Bootstrap support values ≥0.4 are shown at internal nodes. Branch lengths are proportional to the inferred number of amino acid substitutions per site; the scale bar represents 0.05 substitutions per site.

#### 3.2.1. Global D2 backbone and separation of host vs viral lineages

Despite being unrooted, the D2 tree displays a clear partitioning among (i) a broad “representatives” component (including higher plants and structurally characterized algae/cyanobacteria) and (ii) a large marine picocyanobacterial radiation that further subdivides into host-derived lineages and cyanophage-derived lineages. The representative taxa (plants and algae such as *Arabidopsis thaliana*, *Spinacia oleracea*, *Chlamydomonas reinhardtii*, *Chlorella ohadii*, and others) cluster together and provide an external evolutionary reference for interpreting the much denser marine sampling.

A major distinction from the D1 phylogeny is that the D2 tree contains substantially fewer viral sequences, reflecting the lower representation of cyanophage-encoded psbD relative to psbA in our dataset. The cyanophage D2 sequences nevertheless form well-defined, coherent clades corresponding to the *Synechococcus*-associated (Syn.phage) and *Prochlorococcus*-associated (Pro.phage) clades, each supported by strong bootstrap values (0.818 and 0.837, respectively). Together, these patterns suggest that cyanophage *psbD* genes are organized into host-associated reservoirs, albeit with markedly narrower diversity than was observed in the viral *psbA* (D1) dataset described in Sec 3.1 and shown in Fig 1.

#### 3.2.2. *Prochlorococcus* D2 diversification tracks ecotype-level structure

Within the marine portion of the D2 phylogeny, *Prochlorococcus* sequences segregate into well-defined ecotype-specific clusters corresponding to HLI, HLII, LLI, LLII/III, LLIV, LLVII, and LLVIII. Representative strains from these ecotypes (e.g., MED4, MIT9312, NATL1A/NATL2A, MIT9313, and related isolates) occupy distinct positions within the *Prochlorococcus* clade of the tree, consistent with stable, lineage-specific D2 diversification in genomes that encode a single *psbD* homolog.

As in the D1 analysis, the *Prochlorococcus*-centered region of the D2 phylogeny (again referred to as the *Prochlorococcus* clade) includes both host-derived and cyanophage-encoded sequences, forming a single cluster rather than two deeply separated host and viral lineages. The smallest clade containing all *Prochlorococcus* D2 sequences comprises 61 host-derived D2 proteins and 8 cyanophage D2 proteins (total n = 69), similar to the counts in the D1-derived *Prochlorococcus* clade. In contrast to the D1 phylogeny, the basal node defining the *Prochlorococcus* D2 clade receives very strong bootstrap support (0.998), indicating a highly stable evolutionary grouping. Notably, D2 ecotype partitioning is clean and compact at the level of major clades, a pattern expected given that *Prochlorococcus* encodes only a single copy of *psbD* per genome. The absence of paralog ambiguity allows D2 sequence divergence to reflect primarily long-term ecotype differentiation and niche partitioning, rather than isoform turnover or gene duplication, highlighting a more constrained and orderly evolutionary trajectory for D2 compared with D1.

#### 3.2.3. *Synechococcus* D2 forms several clades with limited paralog complexity

The D2 tree contains a broad *Synechococcus* strain sector comprising numerous marine and freshwater *Synechococcus* isolates, as well as several *Synechococcus elongatus* lineages and related taxa (including *Picosynechococcus*). The D2 tree tends to show less paralog-driven expansion than the D1 tree, since *Synechococcus psbA* often occurs in multiple copies (D1.1/D1.2 and additional paralogs) but *psbD* is typically single-copy. This yields a *Synechococcus* D2 phylogeny that is less confounded by isoform interleaving than the D1 tree and may therefore more closely reflect organismal relationships and ecological diversification.

#### 3.2.4. Cyanophage D2 sequences are less abundant than D1 but remain structured by host association

Cyanophage-encoded D2 sequences form two recognizable partitions in the unrooted tree: a large *Synechococcus*-phage sector (containing many “S-” phage names, e.g., S-CAM, S-TIM, and related designations), and a smaller *Prochlorococcus*-phage sector (with “P-” phage names, e.g., P-SSM, P-RSM, P-HM lineages). The existence of distinct Syn-phage vs Pro-phage D2 partitions supports the presence of host-associated viral *psbD* pools, consistent with frequent exchange within host-specific ecological networks. However, *psbD* is retained by a narrower subset of cyanophages, which may reflect stricter functional constraints, a different selective advantage during infection, or a higher loss rate relative to *psbA*.

In summary, the unrooted D2 phylogeny reveals a strong host backbone spanning representative cyanobacteria/algae/plants, and a dense marine picocyanobacterial clade, with ecotype-structured *Prochlorococcus* clustering and a large *Synechococcus* radiation that is less affected by paralogy than D1. Cyanophage D2 sequences form distinct host-associated pools (Syn-phage vs Pro-phage). Still, they are markedly narrower than viral D1, supporting the conclusion that *psbD* retention is less widespread among cyanophages than *psbA*.

### 3.3. Comparative phylogenetic patterns of D1 versus D2 across marine cyanobacteria and cyanophages

Comparison of the D1 (*psbA*) and D2 (*psbD*) phylogenies reveals sharply different *psbA*/*psbD* gene-copy architectures between *Prochlorococcus* and *Synechococcus*. *Prochlorococcus* encodes single-copy *psbA* and *psbD*, and its D1 and D2 sequences thus segregate cleanly by ecotype (HLI/HLII and multiple LL clades). This suggests that both reaction-center subunits track long-term ecotype divergence with minimal paralog confounding. In contrast, *Synechococcus* exhibits extensive *psbA* paralogy with 194 D1 homologs recovered from 166 strains producing multiple D1 types (notably D1.1 and D1.2) that generate a more complex within-host radiation in the D1 tree. The corresponding D2 dataset is substantially less expanded (102 D2 isoforms from 98 strains), reflecting the typical single-copy nature of *psbD* and additional loss or lack of annotation in assemblies. The *Synechococcus* D2 tree topology is thus comparatively more straightforward and more readily interpretable as strain-level diversification rather than isoform turnover.

We also see that the two trees reveal a strong viral gene-repertoire bias, with *psbA* far more broadly retained than *psbD* across cyanophages. *Prochlorococcus* cyanophages frequently encode *psbA*, but only a minority also encode *psbD* (8 of 23 *Prochlorococcus* phages in our dataset), and the same pattern holds for *Synechococcus* phages (72 of 104 carry *psbD*). This disparity is also reflected in the phylogenetic trees, where the D1 tree contains densely sampled phage radiations (both Pro- and Syn-phage), whereas the D2 tree contains more restricted viral radiations. This implies viral D2 is selectively retained by fewer lineages and/or is lost more readily. Importantly, in both D1 and D2, phage sequences nonetheless form recognizable host-associated pools (Syn-phage vs Pro-phage partitions), supporting long-term circulation of viral photosynthesis genes within host-structured ecological networks.

The *Prochlorococcus* ecotype structure aligns with D1/D2 phylogenetic organization. The *Prochlorococcus* ecotype framework, derived from cultivation, physiology, and genomics, aligns closely with the ecotype-structured D1 and D2 phylogenies resolved here, consistent with PSII core protein divergence being associated with long-term light-driven niche differentiation. LL ecotypes (LLI, LLII/III, LLIV, LLVII, and LLVIII) form coherent and clearly distinguishable clades in both trees, consistent with stable lineage divergence in single-copy *psbA* and *psbD* genomes and indicating that PSII evolution in *Prochlorococcus* proceeds primarily through ecotype-level diversification rather than paralog turnover. In contrast, although HLI and HLII ecotypes are ecologically distinct and primarily differentiated by temperature preference, they exhibit minimal divergence in D1 and D2 sequences and consistently cluster as a single HL subclade. This suggests that temperature-associated divergence is not strongly reflected in these D1 and D2 sequences, although temperature effects elsewhere in the genome or physiology cannot be excluded. The clear separation between HL and LL lineages in both phylogenies, together with differences in branching order, highlights systematic protein-level divergence associated with light regime. Given that high irradiance promotes photodamage to PSII reaction-center proteins, these patterns are consistent with a model in which light intensity, rather than temperature, may present a major selective pressure shaping the evolutionary trajectories of D1 and D2 in marine *Prochlorococcus*.

### 3.4. Variations in D1/D2 proteins across *Prochlorococcus* ecotypes and their associated phages

Since the diversification patterns of *Prochlorococcus* D1 and D2 proteins in the phylogenetic trees are largely consistent with their high-light (HL) and low-light (LL) ecotypes, these results suggest that D1 and D2 contain ecotype-associated variation that may reflect light-dependent evolutionary pressures. Accordingly, amino acid positions that consistently differ between HL-and LL-adapted *Prochlorococcus* ecotypes may represent candidate high-light-adaptive substitutions. Comparative sequence alignment revealed pronounced variability in the N-terminal regions of both D1 and D2, with particularly strong divergence between host-derived and phage-encoded sequences. Outside these regions, D1 and D2 proteins are highly conserved across HL and LL *Prochlorococcus* ecotypes and their phage homologs (Supplementary Material). Representative Prochlorococcus strains and cyanophages (Figs. 3 and 4) were selected based on their phylogenetic positions and the similarity of their D1 (PsbA) sequences to the consensus sequence of the corresponding ecotype or cyanophage clade. To capture the overall tree topology, sequences occupying typical, non-outlier positions within the major clades were chosen. Cyanophage representatives were further required to encode both PsbA and PsbD, enabling paired comparisons of phage-encoded D1 and D2 proteins. Despite this overall conservation, numerous ecotype- and phage-associated substitutions were detected throughout both proteins (Figs. 3 and 4). These substitutions distinguish HL- and LL-adapted *Prochlorococcus* strains and are frequently mirrored in phage-encoded homologs. This nonrandom distribution suggests that the observed divergence may be functionally relevant and provides a basis for examining potential differences between HL- and LL-associated D1/D2 proteins.

**Figure 3.**
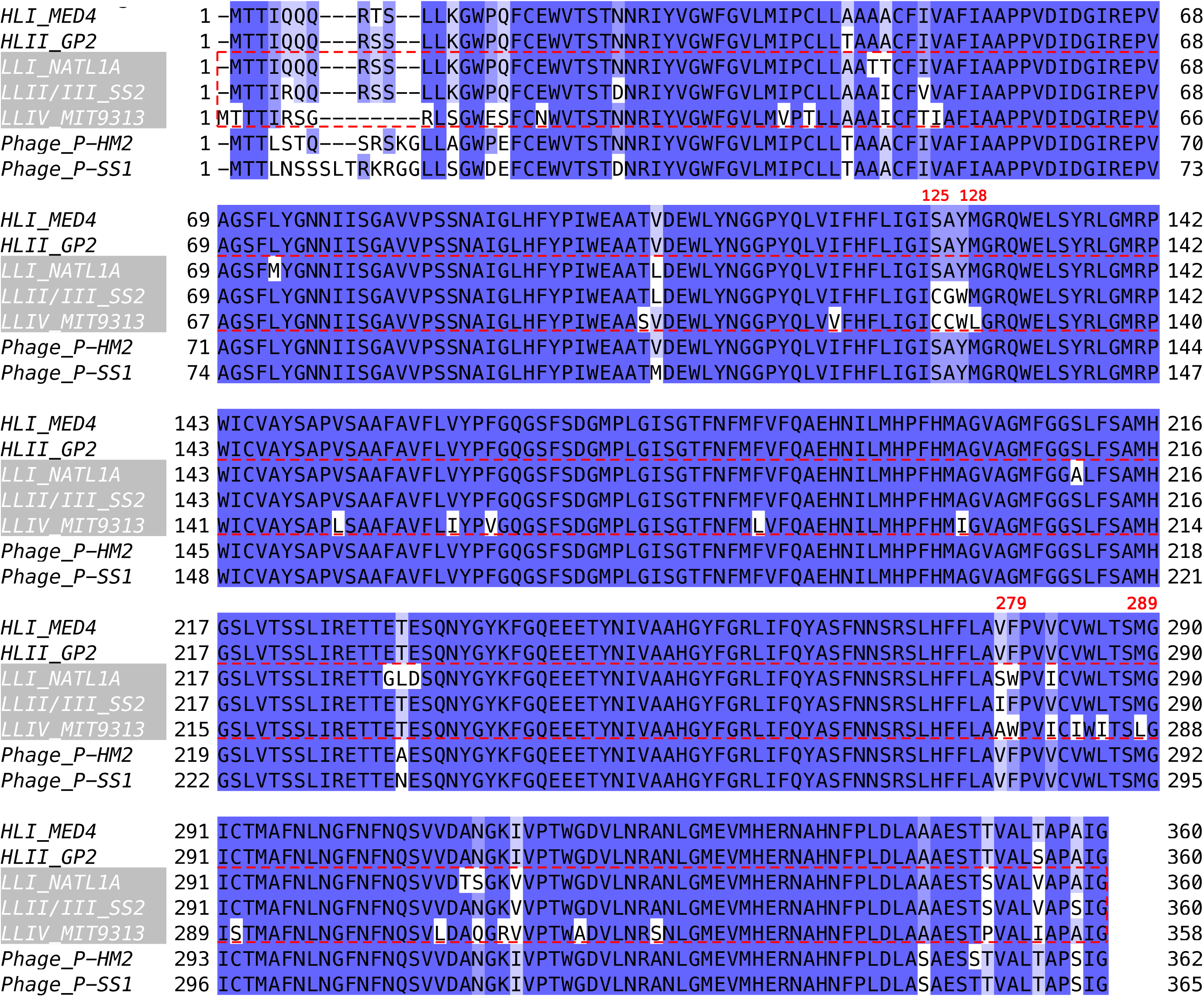
Sequence alignment of PSII D1 (PsbA) protein sequences from representative *Prochlorococcus* strains and cyanophages. MED4 and GP2 represent the high-light-adapted ecotypes HLI and HLII, respectively. NATL1A, SS2, and MIT9313 represent the low-light-adapted ecotypes LLI, LLII/III, and LLIV, respectively. P-HM2 and P-SS1 represent *Prochlorococcus*-infecting cyanophages. Pink arrows in the D1 phylogenetic tree in Fig. 1 indicate the strains and phage sequences analyzed here. Functionally important residues are indicated by bold red numbering. The complete alignment, including all analyzed *Prochlorococcus* strains and phage sequences, is provided in the Supplementary Material. The sequence alignment was generated using Jalview, with residues colored by percentage identity. The shading reflects the proportion of sequences in each column that match the consensus residue: darkest blue (>80% identity), medium blue (60–80% identity), light blue (40–60% identity), and white (<40% identity).

**Figure 4.**
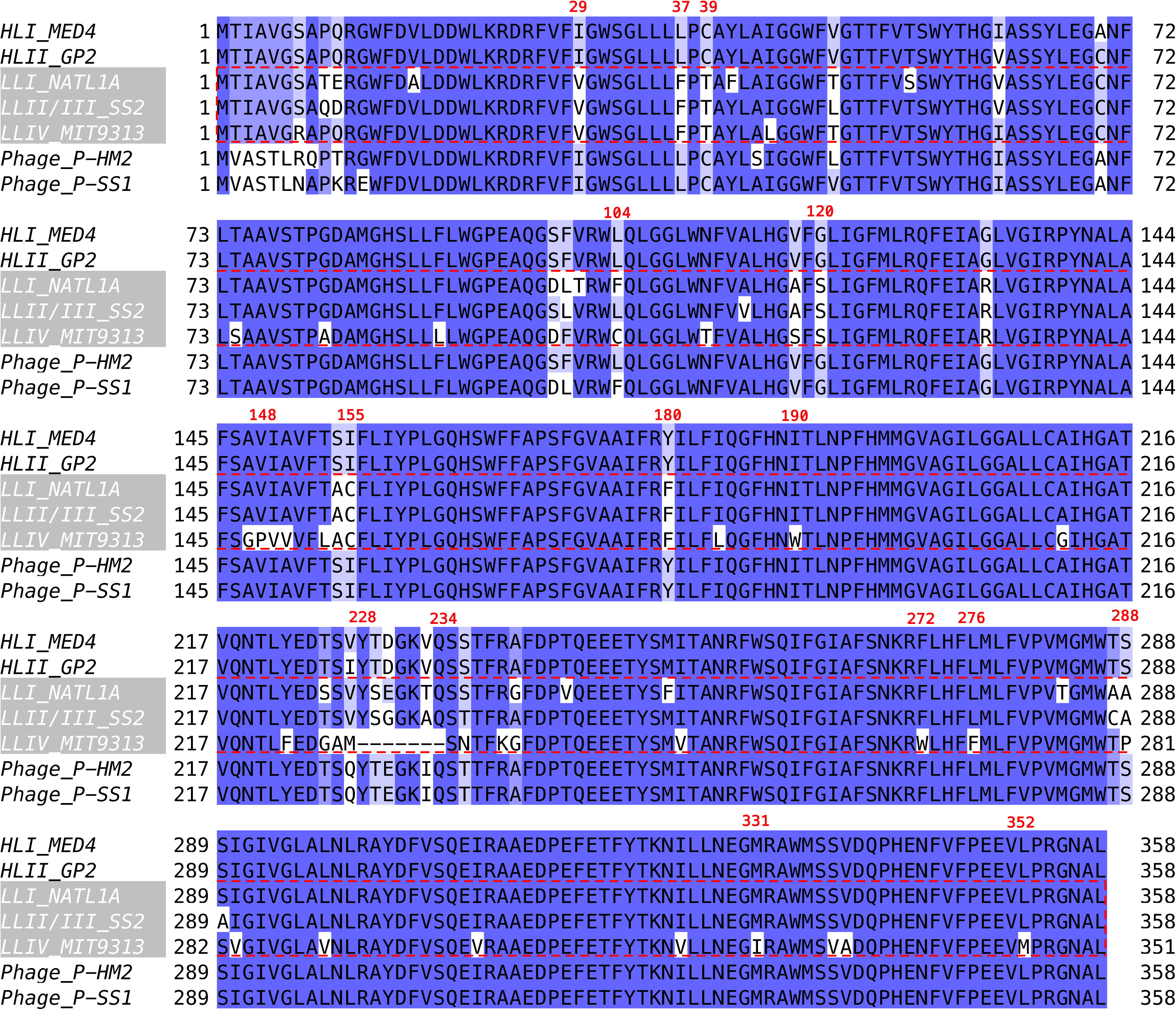
Sequence alignment of the PSII D2 (PsbD) protein from representative *Prochlorococcus* strains and cyanophages. The same strains and phages as in Fig 3 are shown, and coloring and labels are the same as in Fig 3. Pink arrows in the D2 phylogenetic tree in Fig. 2 indicate the representative strains and phage sequences analyzed here.

To provide a structural framework for the functional interpretation of the observed variation in the conserved regions, we used AlphaFold3 to predict the D1/D2/F complex structures of *Prochlorococcus* (*30*). For HL-adapted ecotypes, we used the sequences from *Prochlorococcus* MED4 (HLI ecotype; Fig. 5A), which are highly conserved among HL strains. For LL-adapted ecotypes, we used the corresponding sequences from *Prochlorococcus* MIT9313 (LLIV ecotype; Fig. 5B). Because subunit F is structurally associated with D2, it was included in all predictions. Except for short N-terminal segments, the predicted D1/D2/F complexes exhibited very high confidence scores (pLDDT > 90) across most regions (Figs. 5A, 5B), indicating reliable structural predictions. To further validate these models, we aligned the predicted *Prochlorococcus* complexes to experimentally resolved PSII structures. Among available cryo-EM structures, the D1 and D2 proteins of *Synechocystis* sp. PCC 6803 (PDB: 7N8O; 1.93 Å resolution) showed the highest sequence identity to *Prochlorococcus* MED4 (Table S4) and was therefore used as the reference. Structural alignment using VMD (*31*) showed close correspondence between the predicted and experimental structures, except for the N-terminal regions (Fig. 5C), supporting the reliability of the predicted *Prochlorococcus* D1/D2/F models.

**Figure 5:**
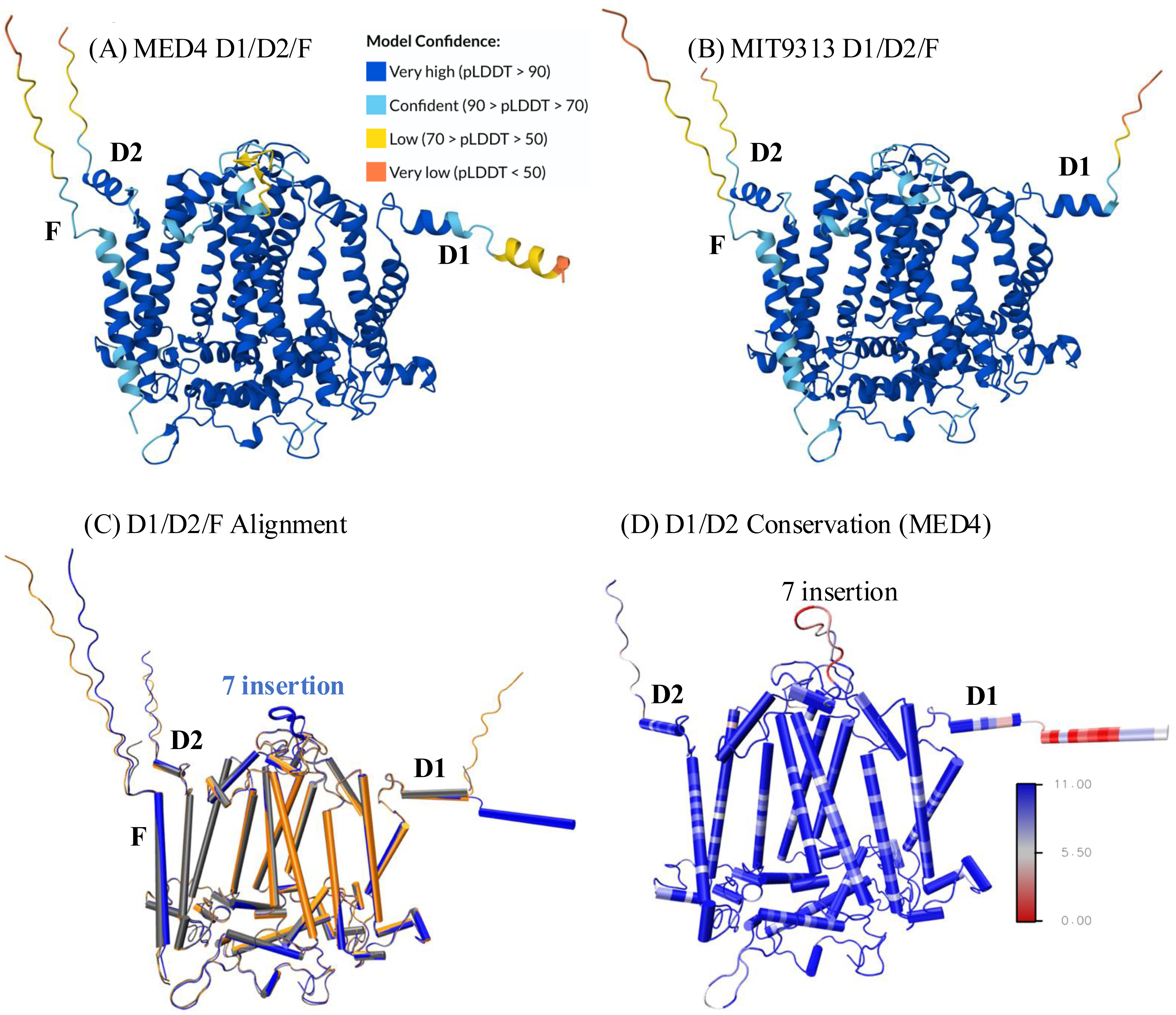
The predicted structures of *Prochlorococcus* D1/D2/F proteins. (**A**, **B**) AlphaFold3-predicted D1/D2/F structures of the *Prochlorococcus* MED4 (HLI) and MIT9313 (LLIV). The D1 protein is the mature form that lacks the C-terminal tail (15 residues). This figure was generated using the AlphaFold3 server. (**C**) Structural alignment of the D1/D2/F proteins from *Prochlorococcus* MED4 (blue), *Prochlorococcus* MIT9313 (orange), and *Synechocystis* 6803 (gray) (PDB: 7N8O). The extended DE loop with a seven-residue insertion in the D2 protein of *Prochlorococcus* MED4 is depicted in the new cartoon representation. (**D**) Residue-level conservation of the D1/D2 proteins. Conservation scores were calculated on the D1/D2 sequences dataset from *Prochlorococcus* strains and their associated phages. Conservation scores were projected onto the AlphaFold3-predicted D1/D2 structure of *Prochlorococcus* MED4. A score of 11 represents complete conservation (blue), whereas a score of 0 indicates no conservation (red).

Residue-level conservation scores calculated in Jalview and projected onto the MED4 D1/D2 structure revealed strong conservation across most of both proteins, interspersed with numerous ecotype-specific variants (Fig. 5D) (*29*, *31*). Notably, the N-terminal region of D1 showed particularly low conservation, largely attributable to the high sequence diversity observed in phage-encoded D1 proteins.

In addition to point substitutions, a comparison of the aligned sequences and structural predictions revealed a seven-residue insertion in the D2 protein, resulting in an extended loop enriched in polar residues. Because this insertion is located between helices D and E, it is hereafter referred to as the extended DE loop. This extended DE loop is present in HL-, LLI-, LLII/III-, and LLVIII-adapted *Prochlorococcus* strains as well as in *Prochlorococcus* cyanophages, but is absent from the LLIV ecotype and from D2 proteins of other oxygenic phototrophs, including *Synechococcus*, cyanobacteria, algae, and plants (Fig. 4). The extended DE loop was observed in both *Prochlorococcus* strains and *Prochlorococcus* cyanophages, suggesting that it may represent a lineage-associated structural feature. Its potential role in PSII function remains to be investigated.

### 3.5. Oxidation-resistant substitutions in D1/D2 proteins across *Prochlorococcus* ecotypes and their associated phages

HL conditions increase the production of reactive oxygen species (ROS) in PSII, potentially leading to oxidative damage (photodamage) to core reaction-center proteins. Variations in oxidation-sensitive residues between HL and LL strains may therefore indicate plausible adaptive mechanisms for enhancing protein stability under HL stress. Based on their intrinsic susceptibility to oxidation, the 20 common amino acids have been classified into four tiers of oxidation sensitivity (Table S3) (*32–34*). Extremely highly oxidation-sensitive residues include cysteine, methionine, tryptophan, tyrosine, and histidine, whereas highly sensitive residues include histidine, phenylalanine, proline, and arginine. Moderately sensitive residues include leucine, isoleucine, valine, lysine, and threonine, while the remaining amino acids are considered low sensitivity (Table S3). Substitutions of highly oxidation-sensitive residues with less oxidation-prone residues are expected to reduce the likelihood of irreversible ROS-mediated side-chain damage in comparable structural contexts. We refer to such substitutions as oxidation-resistant substitutions (ORSs).

Several potential ORSs were identified in the D1 protein (Figs. 3 and 6A; Table 2). At positions 125 and 126 (on helix B in Fig. 6A), cysteine (C) is found in LL-adapted *Prochlorococcus* strains (LLIV and LLII/III). In contrast, serine or alanine is conserved in HL-adapted strains and all *Prochlorococcus* phages. At position 279 (on helix E in Fig 6A), tryptophan (W) is conserved in LLIV and LLI strains but is replaced by phenylalanine (F) in HL-adapted strains and phages. These substitutions reduce intrinsic susceptibility to oxidation while preserving hydrophobic character at those locations.

**Figure 6:**
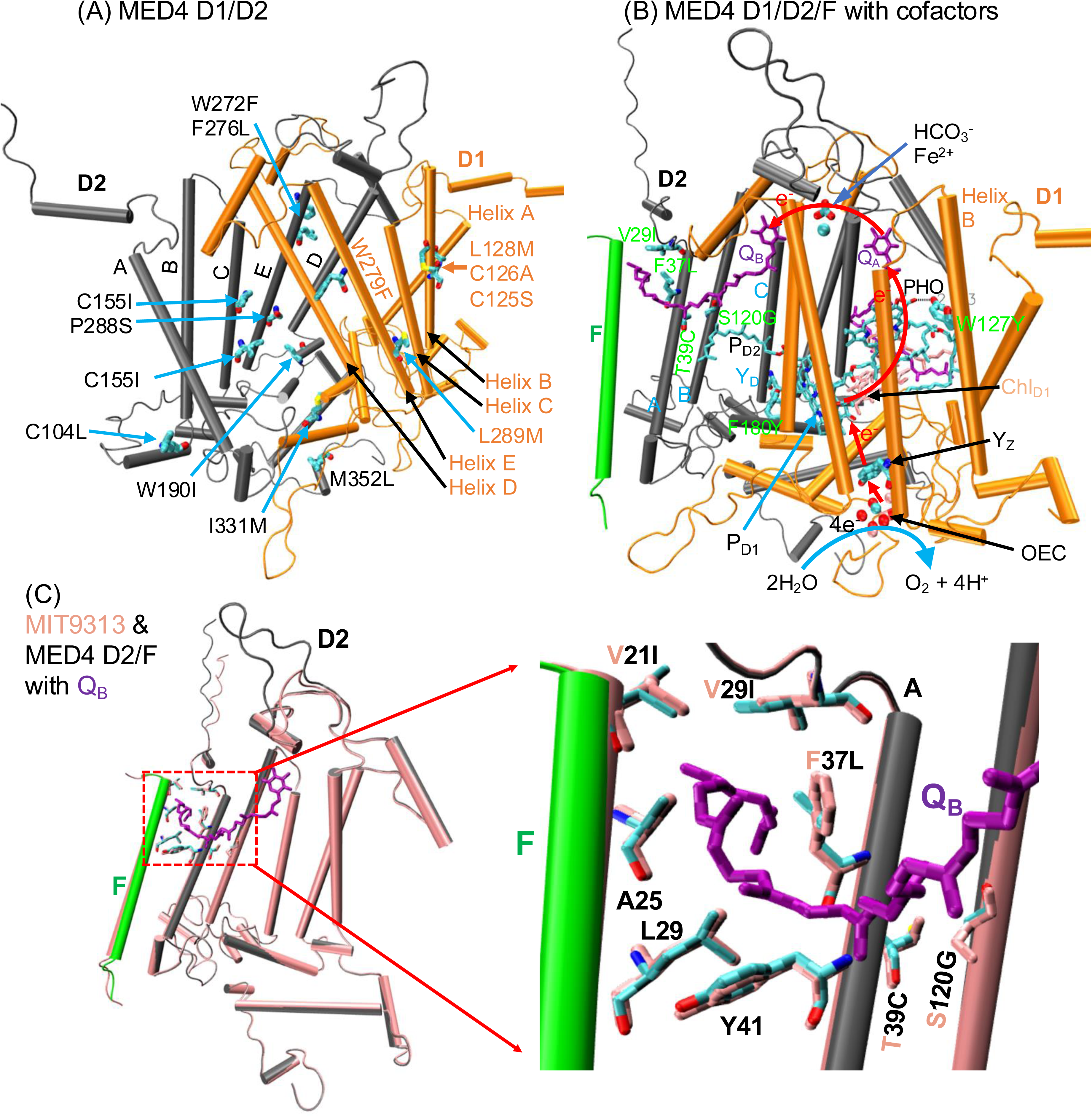
Ecotype-specific variants in *Prochlorococcus* D1/D2 proteins. (A) Oxidation-resistant substitutions in D1/D2 proteins across *Prochlorococcus* ecotypes and their phages. These substitutions were mapped onto the AlphaFold3-predicted D1/D2 structure of *Prochlorococcus* MED4. (B) AlphaFold3-predicted D1/D2 structure of *Prochlorococcus* MED4 with cofactors. The cofactors were incorporated through structural alignment with the cryo-EM structure of PSII from *Synechocystis* (PDB: 7N8O). For clarity, not all cofactors are shown. Only those directly involved in the electron transport chain are displayed, and key residues are labeled. Red arrows indicate the electron transport chain, which begins with a water molecule and proceeds through the Mn cluster, Y_Z_, chlorophyll *a* (P_D1_), chlorophyll *a* (Chl_D1_), phaeophytin (Pheo), plastoquinone (Q_A_), Fe^2+^, and plastoquinone (Q_B_). (C) Structural alignment of the predicted D2/F subunits from *Prochlorococcus* MIT9313 and MED4, representing LLIV and HL ecotypes, respectively. The zoomed-in view highlights residues near Q_B_. The MIT9313 D2/F subunits and corresponding residues are shown in pink.

**Table 2:** Ecotype-specific variants in *Prochlorococcus* D1/D2 proteins and their putative functional effects based on the change from LL to HL, and on structural modeling of substitution locations and cofactors. The residue positions are based on the numbering of the residues in the D1/D2 proteins from *Prochlorococcus* MED4. ORS refers to an Oxidation-Resistant Substitution, RSS refers to an ROS-Suppressing Substitution, and RBS refers to a Redox-Buffering Substitution.

| Protein<br>(No.) | Position | LLIV<br>(15) | LLII/III<br>(10) | LLI<br>(9) | HL<br>(23) | Phage<br>(23/8) * | Putative<br>Effects |
| --- | --- | --- | --- | --- | --- | --- | --- |
| **D1 (80)** | 125 | C | C/S | S | S | S/A | ORS |
|  | 126 | C | A/G | A | A | A | ORS |
|  | 127 | W | W/Y | Y | Y | Y | RSS |
|  | 128 | L | M | M | M | M/A/L | RBS |
|  | 279 | W | F | W | F | F/W | ORS |
|  | 289 | L | M | M | M | M | RBS |
| **D2 (65)** | 29 | V | V | V | I | I | RSS |
|  | 37 | F/L | F | F | L | L | RSS |
|  | 39 | T | T | T | C | C | RSS |
|  | 120 | A/S | S | S | G | G | RSS |
|  | 104 | C | L | F | L | F/L | ORS |
|  | 148 | P | V | V | V | V | ORS |
|  | 155 | C | C | C | I | I | ORS |
|  | 180 | F | F | F | Y | Y | RSS |
|  | 190 | W | I | I | I | I | ORS |
|  | 228-234<br>Insertion | no | yes | yes | yes | yes | Cytosolic<br>binding |
|  | 272 | W | F | F | F | F | ORS |
|  | 276 | F | L | L | L | L | ORS |
|  | 288 | P | A | A | S | S | ORS |
|  | 331 | I | M | M | M | M | RBS |
|  | 352 | M | L | L | L | L | ORS |

Note\*: *Prochlorococcus* phages encode 23 D1 proteins and 8 D2 proteins.

Likely ORSs were also detected in the D2 protein (Figs. 4 and 6A; Table 2). At position 104, cysteine is fully conserved in the LLIV ecotype but is replaced by leucine (L) or phenylalanine in LLII/III, LLI, HL strains, and phages. At position 155, cysteine is conserved in LL-adapted strains, whereas isoleucine is conserved in HL-adapted strains and phages. Similarly, residues P148 and W190 are conserved in the LLIV ecotype but are replaced by V148 and I190, respectively, in other ecotypes and phages. Additional ORSs include W272F, F276L, and M352L, as well as a P288A/S substitution pattern across ecotypes. Collectively, these substitutions reduce the number of oxidation-prone side chains in HL-adapted lineages and phages, consistent with selection for increased resistance to ROS-induced damage.

### 3.6. ROS-suppressing substitutions in D1/D2 proteins across *Prochlorococcus* ecotypes and their associated phages

In addition to reducing susceptibility to oxidation, certain substitutions are expected to suppress ROS generation by modulating PSII electron transfer and charge recombination. In our alignment of D1 proteins, residue 127 is tryptophan in the LLIV and LLII/III ecotypes, but is tyrosine in the LLI and HL ecotypes and in all *Prochlorococcus* phages (Fig. 3; Table 2). Because both residues are highly oxidation-sensitive, this substitution does not constitute an ORS. Structural prediction and alignment revealed a distance of less than 3 Å between the AlphaFold prediction of the phenolic hydroxyl group of Y127 and the O1D atom of pheophytin (Pheo) in the PDB structure of *Synechocystis* (Fig. 6B), suggesting a possible hydrogen bond. As Pheo is a key intermediate in PSII electron transfer, this interaction may stabilize the reduced Pheo□ state and impede charge recombination, thereby reducing the formation of triplet chlorophyll and the subsequent production of singlet oxygen. This analysis suggests that W127Y may act as a ROS-suppressing substitution (RSS): a substitution that reduces primary ROS generation by altering electron-transfer or recombination pathways.

Several ecotype-specific substitutions in the D2 protein may contribute to ROS suppression and adaptation to high-light conditions. At position 29, valine is highly conserved in LL-adapted strains (97%), whereas isoleucine is present in all HL-adapted strains and phages examined (100%). At position 37, phenylalanine, found in 84% of LL-adapted strains, is replaced by leucine in 100% of HL-adapted strains and phages. At position 39, threonine predominates in LL-adapted strains (97%), whereas cysteine is completely conserved in HL-adapted strains and phages (100%). At position 120, serine and alanine occur in 68% and 29% of LL-adapted strains, respectively, but are replaced by glycine in all HL-adapted strains and phages examined. It is worth noting that the F37L substitution reduces side-chain volume by approximately 17.9% and may decrease susceptibility to oxidative modification. The coordinated conservation of the V29I, F37L, T39C, and S/A120G substitutions suggests co-evolutionary adaptation to contrasting light environments. Their complete conservation in HL-adapted strains and phages further indicates that these substitutions may confer a selective advantage under high-light conditions.

Structural analysis placed residues 29, 37, and 39 on the first transmembrane helix, helix A, of the D2 subunit, where they are positioned on or adjacent to the plastoquinone Q_B_ entry/exit channel (Fig. 6B). Structural alignment of the D2/F subunits from *Prochlorococcus* MIT9313 and MED4 further suggests that this channel is predominantly hydrophobic and is shaped by three pairs of residues from D2 helix A and the F subunit. The lower pair, D2-Y41 and F-L29, is conserved in both low-light- and high-light-adapted *Prochlorococcus* strains, suggesting that this region may provide a conserved structural boundary for the channel. The upper pair, D2-V29I and F-V21I, contains substitutions that increase side-chain volume on both sides of the channel. These coordinated changes may shift D2 helix A slightly away from the F subunit, thereby expanding the upper region of the Q_B_ entry/exit pathway.

The D2-F37 and F-A25 residues form the central pair near a putative bottleneck in the Q_B_ entry/exit channel. Because F-A25 is conserved across LL- and HL-adapted *Prochlorococcus* strains, the ecotype-specific difference at this site is primarily associated with the D2-F37L substitution. Phenylalanine is predominant at position 37 in LL-adapted strains. In contrast, leucine is conserved in HL-adapted strains and phages, suggesting an association between this substitution and adaptation to high-light environments. Although AlphaFold3-predicted structures showed only minor differences in static channel width, the altered side-chain chemistry may influence local channel dynamics. The bulky aromatic side chain of phenylalanine could form transient hydrophobic or π-associated contacts with plastoquinone and may impose greater steric constraints near the channel bottleneck. In contrast, leucine lacks an aromatic ring and has a smaller, more conformationally flexible aliphatic side chain, which may reduce local steric constraints and facilitate transient channel opening. These structural considerations suggest that F37L could modulate plastoquinone exchange dynamics even in the absence of a pronounced change in the static channel geometry. Enhanced quinone exchange could, in turn, support more efficient forward electron transfer and reduce the probability of charge recombination under high-light conditions; however, this proposed mechanism requires validation by molecular dynamics simulations and functional experiments.

Together, these substitutions suggest a coordinated remodeling of the Q_B_ entry/exit channel. The V29I substitution may increase side-chain volume and contribute to displacement of D2 helix A away from the F subunit, whereas F37L and T39C may reduce local steric constraints within the channel. Together with S120G, which further reduces side-chain size near the Q_B_-associated region, these changes may facilitate plastoquinone exchange and accelerate forward electron transport. By improving Q_B_ exchange and reducing charge recombination, these substitutions may help limit triplet chlorophyll accumulation and suppress singlet oxygen formation under high-light conditions. We therefore classify V29I, F37L, T39C, and S120G as putative ROS-suppressing substitutions. Further molecular dynamics simulations incorporating PSII cofactors and plastoquinone will be required to test this hypothesis directly.

Finally, phenylalanine is conserved at position 180 of the D2 protein in LL-adapted *Prochlorococcus* strains, whereas tyrosine is conserved at this position in HL-adapted strains and all examined *Prochlorococcus* phages (Fig. 4). Although the F180Y substitution introduces a residue that is generally more susceptible to oxidative modification, structural alignment places residue 180 in the region between the redox-active D2 tyrosine, Y_D_, and the P_D2_ chlorophyll of P680 (Fig. 6B). The proximity of Y180 to this electron-transfer region raises the possibility that its phenolic side chain could influence the local hydrogen-bonding or redox environment. Because oxidized P680 can accept an electron from Y_D_, the introduction of an additional tyrosine near this pathway could potentially provide an alternative or auxiliary redox-active site and thereby alter the kinetics of charge stabilization or recombination. However, the redox activity of Y180 and its ability to participate directly in electron transfer cannot be inferred solely from structural proximity. Thus, F180Y is best considered a candidate redox-modulating substitution associated with high-light adaptation, rather than an oxidation-resistant substitution.

### 3.7. Distribution of redox-buffering residues in D1 and D2 proteins across *Prochlorococcus* ecotypes and their associated cyanophages

Methionine and cysteine are among the most oxidation-sensitive amino acids. They can function as redox-buffering residues by scavenging reactive oxygen species (ROS) generated within Photosystem II (PSII), particularly under HL conditions (*35*, *36*). To evaluate whether the abundance and distribution of these residues vary across *Prochlorococcus* ecotypes and their infecting cyanophages, we quantified the abundance of methionine (Met) and cysteine (Cys) residues in the D1 and D2 subunits (Table 3).

**Table 3:** Counts of methionine and cysteine residues in Photosystem II D1/D2 proteins across *Prochlorococcus* ecotypes and their infecting cyanophages.

| Strain/Phage | Methionine |  |  | Cysteine |  |  |
| --- | --- | --- | --- | --- | --- | --- |
|  | D1 | D2 | D1+D2 | D1 | D2 | D1+D2 |
| HL | 14 | 11 | 25 | 6 | 2.9 | 8.9 |
| LLI | 15 | 9 | 24 | 6.1 | 3 | 9.1 |
| LLII/III | 14 | 11 | 25 | 6.7 | 4.2 | 10.9 |
| LLIV | 12 | 12 | 24 | 6 | 4 | 10 |
| LLVII | 15 | 12 | 27 | 7.3 | 3 | 10.3 |
| LLVIII | 14 | 9 | 23 | 7 | 4 | 11 |
| Phages | 13.7 | 11.1 | 24.8* | 5 | 2.1 | 7.1* |

\*: Most *Prochlorococcus* phages encode only the PSII D1 protein and not D2. Therefore, the total counts of methionine and cysteine residues in phage-encoded D1/D2 proteins are provided solely as a reference.

In the D1 subunit, the LLIV ecotype contains twelve methionine residues, whereas other LL and HL *Prochlorococcus* ecotypes (as well as *Prochlorococcus* phages) generally contain a higher number of methionine residues. In contrast, the LLIV ecotype contains twelve methionine residues in the D2 subunit, while other LL and HL ecotypes and phages contain fewer methionine residues. Despite this redistribution between subunits, the combined number of methionine residues across D1 and D2 remains broadly similar among *Prochlorococcus* ecotypes and their associated cyanophages (Table 3), indicating conservation of overall methionine-based redox-buffering capacity at the level of the PSII reaction center.

Structural mapping of methionine residues onto the predicted *Prochlorococcus* PSII structure shows that a substantial fraction is positioned in proximity to major redox-active cofactors, including chlorophyll *a* molecules (P_D1_/P_D2_ and Chl_D1_/Chl_D2_), pheophytin (Pheo), plastoquinone (Q_A_/Q_B_), and the oxygen-evolving complex (OEC), as shown in Fig. 7A. These regions are well-known to be centers of ROS formation during charge recombination and electron leakage under HL conditions. The apparently non-random spatial enrichment of methionine residues near these cofactors is consistent with a role in localized redox buffering, whereby reversible methionine oxidation could mitigate oxidative damage to nearby photochemical components. We note while the ORS and RSS substitutions may confer protection by reducing oxidative stresses near the electron transport chain, an enhancement of ROS-scavenging or redox-buffering capacity arising from the gain or retention of the oxidation-sensitive Met and Cys may confer protection through a different mechanism.

**Figure 7.**
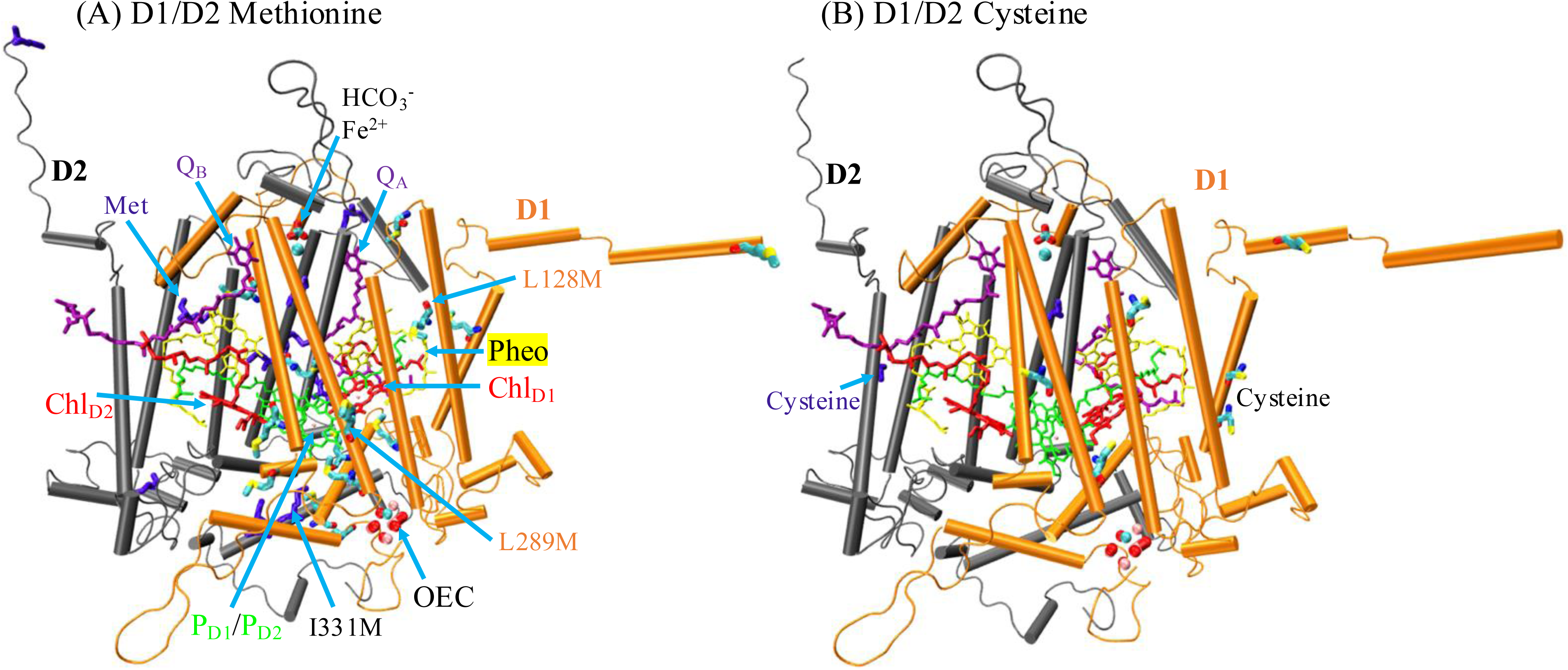
Spatial distribution of methionine and cysteine residues in Photosystem II D1 and D2 subunits of *Prochlorococcus* MED4. (A) Methionine residues are mapped onto the PSII D1 and D2 subunits. Chlorophyll *a* molecules (P_D1_/P_D2_ and Chl_D1_/Chl_D2_) are shown in green and red, respectively; pheophytin (Pheo) and plastoquinone (Q_A_/Q_B_) are shown in yellow and purple, respectively. Methionine residues in the D1 subunit are colored cyan, whereas those in the D2 subunit are colored violet. Key mutants involving redox-buffer substitutions (RBS) are indicated. (B) Cysteine residues mapped onto the PSII D1 and D2 subunits. Cofactors are shown using the same color scheme as in panel (A).

Based on this structural context, we define redox-buffering substitutions (RBS) as amino-acid replacements that introduce methionine residues at positions previously occupied by non–redox-active side chains, potentially enhancing the local capacity for ROS scavenging within PSII. In the D1 protein, residues L128 and L289 are conserved as leucine in the LL–adapted *Prochlorococcus* LLIV ecotype but are substituted by methionine in other *Prochlorococcus* ecotypes and in cyanophage-encoded D1 proteins (Table 2). In the D2 protein, residue I331 shows a similar pattern, being conserved in LLIV but replaced by methionine in other *Prochlorococcus* ecotypes and associated cyanophages (Table 2).

Structural mapping localizes the D1 L128M substitution to the vicinity of pheophytin. The D1 L289M and D2 I331M substitutions are near (within 1nm) of the primary chlorophyll *a* (P_D1_/P_D2_) (Fig. 7A). Given their proximity to key redox cofactors and known ROS-generation sites, these substitutions introduce methionine residues at strategically exposed positions within the reaction center. We therefore classify L128M and L289M as D1 redox-buffering substitutions and I331M as a D2 redox-buffering substitution, all of which likely contribute to enhanced oxidative resilience of PSII under HL conditions.

Cysteine residues display a more ecotype-independent distribution. Across *Prochlorococcus* ecotypes, the D1 subunit contains approximately six cysteine residues, while the D2 subunit contains approximately four, yielding a total of ∼10 cysteine residues per PSII reaction center. This total is notably higher than that observed in *Prochlorococcus* cyanophages, which average approximately 7 cysteine residues per D1/D2 pair (Table 3). Structural mapping shows that cysteine residues in both D1 and D2 are located near (within 1nm of) key cofactors (similar to the methionine locations described above), including chlorophyll *a*, pheophytin, and plastoquinone, as shown in Fig. 7B. This positions them to intercept ROS produced during electron transfer reactions and act as a redox buffer.

Together, these results indicate that redox-buffering residues are strategically distributed within PSII rather than randomly positioned. Although the total number of methionine and cysteine residues is broadly conserved across *Prochlorococcus* ecotypes, their redistribution between D1 and D2 subunits and their reduced abundance in phage-encoded PSII proteins suggest differential optimization of redox buffering between hosts and phages. This organization supports a model in which *Prochlorococcus* maintains an intrinsic, protein-encoded redox-buffering system to mitigate photodamage under persistent HL stress. At the same time, cyanophages retain a streamlined yet host-compatible subset of these protective features.

## 4. Discussion

### 4.1. Ecotype-structured PSII evolution in *Prochlorococcus*

Our analyses reveal that both D1 and D2 proteins in *Prochlorococcus* vary in close correspondence with ecotype structure, with the LL and HL lineages forming coherent, well-supported phylogenetic groupings. Because each *Prochlorococcus* genome encodes a single copy of *psbA* and *psbD*, this structure is not confounded by paralog turnover. Instead, it is consistent with long-term lineage divergence driven by ecological niche specialization (*1–3*). The clear segregation of LL ecotypes (LLI, LLII/III, LLIV, LLVII, and LLVIII) and the tight clustering of HLI and HLII into a single HL subclade suggest that light intensity (rather than temperature) may contribute to divergence in PSII core proteins (*6*, *9*, *37*). Notably, the placement of LLVII and LLVIII as distinct yet evolutionarily intermediate lineages is consistent with recent cultivation-based evidence indicating that these ecotypes occupy upper- to mid-euphotic niches and exhibit mixed physiological traits, thereby reinforcing the biological relevance of their phylogenetic positions (*4*).

Comparison of the D1 and D2 phylogenetic trees revealed distinct patterns of sequence diversity and clustering (Figs. 1 and 2). The D1 tree displayed greater topological complexity, particularly among *Synechococcus* and cyanophage sequences, with multiple closely related clusters and a broader representation of viral sequences. In contrast, the D2 tree was more compact, contained fewer cyanophage sequences, and showed less extensive branching among the sampled lineages. Both trees retained broad separation among major host groups and *Prochlorococcus* ecotypes, although these patterns were more complex in D1. The greater heterogeneity observed in the D1 tree is consistent with the presence of multiple *psbA* isoforms in some *Synechococcus* genomes and the more frequent occurrence of *psbA* than *psbD* in cyanophages. Thus, within the present dataset, D1 exhibits greater sequence and topological diversity than D2.

A limitation is that phylogenetic clustering and structure-based analyses can identify candidate adaptive substitutions, but they cannot directly demonstrate that these substitutions were driven by HL selection. Establishing such evolutionary pressure would require analyses that distinguish selection from shared ancestry, recombination, horizontal gene transfer, and neutral divergence. These could include ancestral-state reconstruction of HL and LL transitions, mapping candidate substitutions across independent ecotype transitions, codon- or protein-level tests for episodic or diversifying selection, recombination-aware gene-tree inference, and population-genomic comparisons along environmental light gradients. Functional validation would also be needed, for example, by introducing candidate HL substitutions into the LL ecotype, or reversing them in the HL ecotype, followed by measurements of PSII activity, photodamage, ROS production, repair kinetics, and fitness under controlled HL and LL conditions. Thus, the substitutions identified here are best interpreted as structure-informed, HL-associated candidates that generate testable hypotheses about adaptation, rather than as direct evidence of selection.

### 4.2. Cyanophage D1/D2 evolution remains host-coupled

Rather than forming deeply diverged viral lineages, cyanophage-encoded D1 and D2 sequences cluster within host-associated diversity, especially within the *Prochlorococcus* clade. This pattern supports a model in which viral *psbA* and *psbD* genes are repeatedly acquired from hosts and subsequently homogenized through recombination, rather than evolving independently (*20*, *21*). The preferential clustering of *Prochlorococcus* phage D2 sequences near HL host clades further suggests that viral PSII components are selectively optimized for HL environments, consistent with experimental observations that phage infection can maintain or enhance host photochemical performance under HL conditions (*20*, *24*). These findings reinforce the view that cyanophages function as dynamic reservoirs of photosynthetic genes that both reflect and reinforce host ecological adaptation (*38*).

The higher prevalence of cyanophage-encoded *psbA* relative to *psbD* in our dataset is unlikely to reflect sampling bias; rather, it likely reflects distinct functional constraints and turnover rates of the D1 and D2 reaction-center proteins. D1 is the primary photodamage-sensitive subunit of PSII and is rapidly degraded and replaced during the PSII repair cycle, particularly under high-light stress. Thus, retention of phage-encoded *psbA* may confer an adaptive advantage by enabling infected host cells to maintain PSII activity and sustain photosynthetic electron transport during viral replication. By contrast, D2 is generally more structurally stable and replaced less frequently, resulting in weaker selective pressure for cyanophages to retain *psbD*. This difference in selective demand may explain why phage-encoded D2 proteins are less common than phage-encoded D1 proteins

### 4.3. Multiple protein-level strategies mitigate HL-induced photodamage

By integrating sequence variation with structural context, we identify several complementary mechanisms by which D1/D2 proteins may mitigate HL-induced photodamage. Oxidation-resistant substitutions (ORS) reduce the prevalence of highly oxidation-sensitive residues at structurally vulnerable positions, lowering the likelihood of irreversible side-chain damage (*11*, *12*, *14*). ROS-suppressing substitutions alter the local steric and electrostatic environments near redox-active cofactors, plausibly modulating charge recombination pathways and limiting singlet oxygen formation (*13*, *16*, *17*). In parallel, the enrichment and spatial distribution of methionine and cysteine residues near chlorophylls, pheophytins, quinones, and the oxygen-evolving complex suggest an additional redox-buffering layer, whereby reversible oxidation of sulfur-containing residues scavenges ROS before critical functional groups are damaged (*18*, *19*).

Notably, methionine and cysteine may contribute to two mechanistically distinct protective strategies. As oxidation-sensitive residues, Met and Cys can represent vulnerable sites for oxidative modification; therefore, their loss at structurally exposed or functionally sensitive positions may improve protein oxidation resistance by reducing potential targets of irreversible oxidative damage. Conversely, the gain or retention of Met and Cys near redox-active cofactors may protect PSII through a different mechanism, by providing local ROS-scavenging or redox-buffering capacity. Thus, these two strategies should be considered separately: one reduces oxidation-sensitive targets, whereas the other may enhance the ability of D1/D2 proteins to buffer ROS generated during high-light stress.

The observation that D1 consistently contains more methionine residues than D2 across most *Prochlorococcus* ecotypes and their phages aligns with the well-established role of D1 as the primary photodamage and turnover target in PSII under HL stress (*12*, *14*). This asymmetry supports a model in which redox buffering is preferentially concentrated in the subunit most exposed to ROS-generating reactions, enhancing system-level robustness without compromising core photochemistry.

### 4.4. Implications for the evolution of oxygenic photosynthesis

Taken together, our results highlight how subtle, distributed amino-acid changes that do not appear to alter the structure of the D1/D2 complex significantly may contribute to PSII function under chronic HL stress. The association of ecotype-specific substitutions and cyanophage-host coupled gene pool underscores that PSII evolution in marine environments should be interpreted in the context of viral gene flow and selection (*20*, *38*, *39*). More broadly, this work illustrates how combining phylogenetics with structure-aware analyses can reveal mechanistic hypotheses about the diversification of one of Earth’s most consequential molecular machines (*17*).

### 4.5. Summary

This study provides a comprehensive, structure-aware view of HL adaptation in PSII across marine picocyanobacteria and their cyanophages. By integrating phylogenetic reconstruction, sequence analysis, and AlphaFold3-based structural modeling, we demonstrate that D1 and D2 sequence variation in *Prochlorococcus* is tightly linked to ecotype differentiation and that cyanophage-encoded reaction-center proteins cluster with host-associated diversity. We identify multiple, complementary protein-level strategies—including oxidation-resistant substitutions, ROS-suppressing substitutions, and redox-buffering residue distributions—that may collectively enhance PSII resilience under HL stress. These findings emphasize the potential role of fine-scale protein evolution and viral–host gene exchange in shaping the performance and durability of oxygenic photosynthesis in the modern ocean, offering new mechanistic insights into how photosynthetic systems adapt to extreme and persistent light environments.

## CRediT authorship contribution statement

**Rulong Ma** Conceptualization; Methodology; Software; Validation; Formal analysis; Investigation; Data curation; Writing - original draft; Writing-review & editing; Visualization. **Ruonan Wu** Writing-review & editing. **Greg Morrison**: Resources, Writing-Review & Editing, Supervision, Project administration, Funding acquisition.

## Acknowledgements

This work was completed in part with resources provided by the Research Computing Data Core at the University of Houston. We gratefully acknowledge Drs. Margaret Cheung and Amity Andersen of Pacific Northwest National Laboratory (PNNL) for their coordination and support of this collaboration.

