## Supplementary Material for "Molecular Basis of High-light Adaptation in Cyanobacteria and Cyanophages through the D1/D2 subunits of Photosystem II"

**Supplemental material**

**Supplemental Table S1**: Summary of *Prochlorococcus* strains

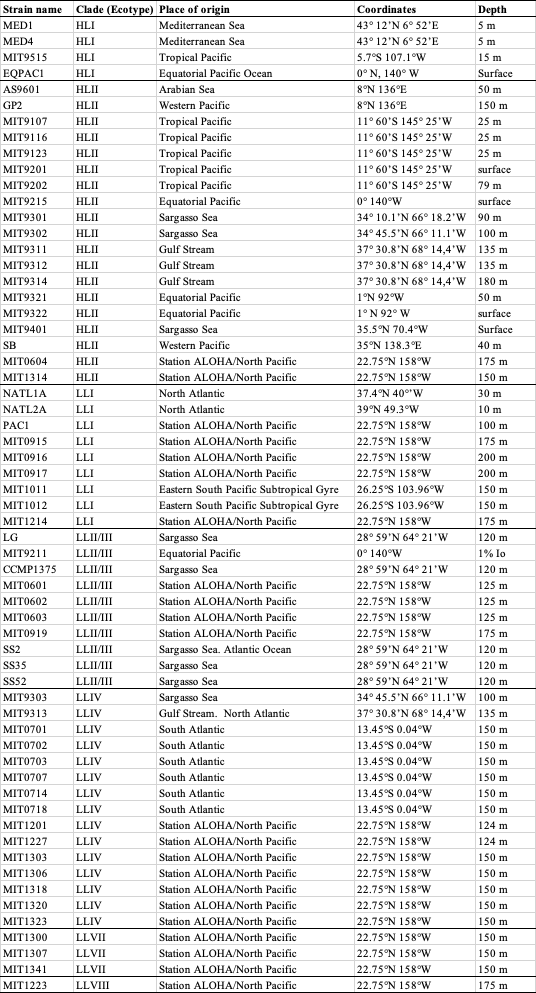

Note: 1% Io = the depth where only 1% of surface light remains after attenuation by water. All data in this table are derived from the Chisholm Lab (Massachusetts Institute of Technology) Prochlorococcus stock cultures resource (https://chisholmlab.mit.edu/cultures/prochlorococcus-stock-cultures/).

**Supplemental Table S2**: Summary of cyanobacteria–cyanophage host–virus relationships*.

| **Host** | **Host strain** | **Phage** |
| --- | --- | --- |
| ***Prochlorococcus*** | **MED4** | P-HM2, MED4-213, P-HM1, P-SSM4, P-SSP7, P-GSP1 |
|  | **MIT9515** | 9515-10a, P-TIM68 |
|  | MIT9312 | P-SSP2, P-SSP3 |
|  | NATL1A | NATL1A-7, P-SSM2, P-SSM7 |
|  | NATL2A | P-RSM3, P-SSM3, NATL2A-133, P-RSM6, P-SSM5, P-SSP10, P-TIM40 |
|  | MIT9313 | P-SS1 |
|  | MIT9303 | P-RSM1, P-RSM4 |
| ***Synechococcus*** | WH7803 | S-PM2, S-CAM1, S-CAM3, S-CAM8, S-CAM9, S-IOM18, S-RIM2, S-RIM8, S-SKS1, KBS-M-1A, S-RIP2, KBS-P-1A, Syn30, Syn33 |
|  | WH8103 | S-RSM4 |
|  | WH8102 | S-ShM2, S-SSM2, S-SSM5, S-TIM5, Syn1, S-TIM4 |
|  | WH6501 | S-SM1 |
|  | WH8107 | S-SM2, Syn10 |
|  | WH8109 | Syn19, S-SSM6a, S-SSM6b, S-TIP37 |
|  | WH8012 | Syn2, Syn9, S-SSM7 |
|  | CB0101 | S-CBM2, S-CBP3, S-CBP4 |
|  | LC16 | S-CRM01 |
|  | WH8101 | S-RIP1 |
|  | WH8018 | S-SSM4 |
|  | Unknow | S-MbCM6, S-MbCM6, S-RIM44, S-RIM50, S-WAM2, KBS-P-1A, metaG-MbCM1 |

*Note: each cyanophage carries only the *psbA* gene or *psbA*/*psbD* genes. This data comes from a previous review paper (*1*).

**Supplemental Table S3**: Classification of the 20 common Amino Acids by Oxidation Susceptibility (*2*–*6*)

| **Oxidation Tier** | **Amino Acid** | **1-letter** | **Relative Susceptibility** | **Reversibility** |
| --- | --- | --- | --- | --- |
| **Tier 1: Extremely high** | Cysteine | C | Very high | Partial |
|  | Methionine | M | Very high | Yes |
|  | Tryptophan | W | Very high | No |
|  | Tyrosine | Y | Very high | No |
| **Tier 2: High** | Histidine | H | High | No |
|  | Phenylalanine | F | High | No |
|  | Proline | P | High | No |
|  | Arginine | R | High | No |
| **Tier 3: Moderate** | Leucine | L | Moderate | No |
|  | Isoleucine | I | Moderate | No |
|  | Valine | V | Moderate | No |
|  | Lysine | K | Moderate | No |
|  | Threonine | T | Moderate | No |
| **Tier 4: Low** | Alanine | A | Low | No |
|  | Glycine | G | Low | No |
|  | Serine | S | Low | No |
|  | Asparagine | N | Low | No |
|  | Glutamine | Q | Low | No |
|  | Aspartate | D | Very low | No |
|  | Glutamate | E | Very low | No |

**Supplemental Table S4**: Sequence identity of PSII D1/D2 proteins between *Prochlorococcus* strain MED4 and organisms with experimentally determined PSII structures

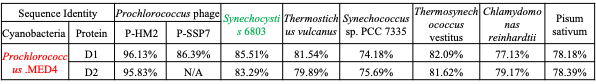
